# miR-128-induced LINE-1 restriction is dependent on down-regulation of hnRNPA1

**DOI:** 10.1101/195560

**Authors:** Lianna Fung, Herlinda Guzman, Evgueni Sevrioukov, Adam Idica, Eddie Park, Aurore Bochnakien, Iben Daugaard, Douglas Jury, Ali Mortazavi, Dimitrios G Zisoulis, Irene Munk Pedersen

## Abstract

The majority of the human genome is made of transposable elements, giving rise to interspaced repeats, including Long Interspersed Element-1s (LINE-1s or L1s). L1s are active human DNA parasites involved in genomic diversity and evolution, but can also contribute to genomic instability and diseases. L1s require host factors to complete their life cycles, whereas the host has evolved numerous mechanisms to restrict L1-induced mutagenesis. Restriction mechanisms in somatic cells include methylation of the L1 promoter, anti-viral factors and RNA-mediated processes such as small RNAs. microRNAs (miRNAs or miRs) are small non-coding RNAs that post-transcriptionally repress multiple target genes often found in the same cellular pathways. We have recently established that the interferon-inducible miR-128 function as a novel restriction factor inhibiting L1 mobilization in somatic cells. We have further demonstrated that miR-128 function through a dual mechanism; by directly targeting L1 RNA for degradation and indirectly by inhibiting a cellular co-factor which L1 is dependent on to transpose to new genomic locations (TNPO1). Here we add another piece to the puzzle of the enigmatic L1 life cycle. We show that miR-128 also inhibits another key cellular factor, hnRNPA1, by significantly reducing mRNA and protein levels through direct interaction with the coding sequence (CDS) of hnRNPA1 mRNA. Furthermore, we demonstrate that repression of hnRNPA1 using shRNA significantly decreases *de novo* L1 retrotransposition and that induced hnRNPA1 expression enhances L1 mobilization. Finally, we determine that hnRNPA1 is a functional target of miR-128 and that induced hnRNPA1 expression in miR-128-overexpressing cells can partly rescue the miR-128-induced repression of L1’s ability to transpose to different genomic locations. Thus, we have identified an additional mechanism by which miR-128 represses L1 retrotransposition and mediate genomic stability.

## INTRODUCTION

Long-interspaced nuclear elements-1 (LINE-1 or L1) are the only autonomous transposable elements and account for approximately 17% of the human genome [1-3]. Although the majority of L1 encoded in the genome is truncated and inactive, intact and active L1 is ∼6 kilobase pairs (kb) in length and contains a 5’UTR, three open-reading frames – ORF1, ORF2 and ORF0 – and a short 3’UTR. The 5’UTR has promoter activity in both the sense and antisense direction [4-6]. ORF1 encodes a 40 kDa protein with RNA-binding and nucleic acid chaperone activity and ORF2 encodes a 150 kDa protein with endonuclease and reverse transcriptase activities [7-9]. ORF0 is transcribed in the antisense direction and encodes a protein that enhances L1 retrotransposition, however the exact mechanism remains unknown [10]. L1 mobilizes and replicates via an RNA by a “copy and paste” mechanism via an RNA intermediate [11, 12]. Integration of L1 at new locations in the genome generates mutations that can create new genes or affect gene expression [13, 14]. L1 retrotransposition is associated with a variety of diseases ranging from Hemophilia to cancer and developmental abnormalities [2, 15-18].

As a result, multiple mechanisms have evolved to regulate L1 activity. In germ cells, specific small RNA subtypes (piRNAs) efficiently counteract L1 activity [19, 20]. In somatic cells, L1 transcription is inhibited by DNA methylation of the L1 promoter [21, 22]. In hypomethylated cell populations such as cancer cells or pluripotent stem cells, the L1 promoter is often de-repressed allowing for active L1 replication retrotransposition [22-24]. Under these conditions other mechanisms of L1 restriction are important, including suppression by DNA and RNA editing proteins, such as AID, APOBECs and ADAR [25, 26], as well as the microprocessor [27].

The majority of the human transcriptome is believed to be under miR regulation, suggesting that the post-transcriptional control of gene regulation by non-coding RNA (ncRNA) may rival that of proteins in regulating multiple genetic pathways [28, 29]. miRs are endogenously encoded 21–24-nucleotide (nt) RNAs that regulate the expression of mRNAs containing complementary sequences. After transcription and processing in the nucleus, the mature miR is loaded onto specific Argonaute (Ago) proteins - referred to as a miR-induced silencing complex (miRISC) – in the cytoplasm. The miRISC then binds to partially complementary mRNA sequences and mediates mRNA degradation and translational [29, 30]. Complementarity between the miR (5’ position 2-7) and a mRNA target “seed” site usually results in reduced protein expression through a variety of mechanisms that involve mRNA degradation and translational repression [30, 31].

We recently established that miR-128 represses the activity of L1 retrotransposons in somatic cells. We described a novel mechanism for this regulation, by which miR-128 binds directly to the ORF2 coding region sequence of L1 RNA resulting in L1 repression [32]. In addition, we have recently expanded the mechanistic studies of miR-128-induced L1 restriction, demonstrating that miR-128 also represses the cellular nuclear import factor Transportin 1 (TNPO1), which we find L1 is dependent on for *de novo* retrotransposition (Idica et al. *in revisions* 2017). It is well established that miRs repress multiple cellular mRNAs by binding to homologous target seed sequences; the proteins of these target mRNAs frequently function in the same pathway, suggesting that miRs act to fine-tune specific cellular networks [33-36].

In this study, we wished to identify additional cellular targets of miR-128 that may be involved in the L1 retrotransposition pathway. Here we report that miR-128 also represses retrotransposition by targeting a key cellular factor required for L1 retrotransposition, namely hnRNPA1 (heterogenous nuclear ribonucleoprotein A1). hnRNPA1 is one of the most abundant proteins in the nucleus and is expressed in all cell types and tissues [37]. The hnRNPA1 proteins is involved in both DNA and RNA metabolism including genomic stability and telomere binding [38, 39]. hnRNPA1 is bound to poly (A) sequences of RNA on both the cytoplasm and nucleus [40] and accompanies mature transcripts through the Nuclear Pore Complex, supporting its proposed role in mRNA shuttling [41]. The nuclear localization of hnRNPA1 depends on a 38aa long nuclear localization sequence known as M9 [42-44], which binds to TNPO1 to mediate shuttling through the nuclear pore complex [39, 45-49]. It has been described that hnRNPA1 interact with L1 ORF1p through an RNA bridge, likely as part of the L1-RNP complex [50]. In this study, we show that miR-128 regulates hnRNPA1 levels and demonstrate that hnRNPA1 is required for efficient L1 replication and *de novo* retrotransposition. Thus, we have discovered another key player in the L1 life cycle, which is subjected to miR-128 regulation.

## RESULTS

### Identification of miR-128 targets which function as co-factors for L1 retrotransposition

We have previously demonstrated that miR-128 directly targets L1 RNA and represses *de novo* retrotransposition and genomic integration in somatic cells [32]. Furthermore, we recently determined that miR-128 also regulates cellular co-factors, some of which L1 may be dependent on. It has previously been debated whether L1-RNP is only dependent on cell division to access host DNA [51-53]. However, the hypothesis could not account for how L1 integrates back into the genome in non-dividing cells such as neurons [54]. For this reason we were excited to determine that miR-128 also targets the nuclear import factor Transportin 1 (TNPO1), resulting in reduced nuclear import of L1 Ribonuclear Protein (L1-RNP) complexes (Idica et al. *in revisions*). However as TNPO1 does not contain a nuclear localization signal (NLS), many questions were left unanswered.

In order to identify additional miR-128 targets involved in the regulation of L1 retrotransposition, and add to our understanding of the L1 life cycle, we performed a screen for potential miR-128 targets utilizing DGCR8^-/-^ mouse embryonic stem cells (mESCs) (a kind gift from Dr. Blelloch). DGCR8 is a critical component of the microprocessor involved in processing pri-miRs into their mature forms [55]. In DGCR8^-/-^ mESC cells, pri-miR transcripts cannot be further processed by the microprocessor and are not loaded into a RISC complex [56]. As a result, the DGCR8^-/-^ system is free of mature, biologically active canonical miRs. We transfected DGCR8^-/-^ mESCs with miR controls or miR-128 in triplicate cultures and harvested cells after 12hours in order to enrich for primary target mRNAs, as opposed to studying secondary effects of miR-128. Two replicates of each triplicate were selected and cDNA libraries were generated using the Smart-seq2 protocol [57]. The libraries were sequenced as 43 bp paired-end reads. STAR was used to align the reads on to the mm9 genome [58]. RSEM [59] was used to quantitate the gene expression and EBSeq [60] was used to identify differentially expressed genes. We then performed overlay analysis of the identified miR-128 targets with previously reported results from a proteomic screen identifying L1 ORF1p-encoded protein (ORF1p) interaction partners [50] (see Figure 1A). Interestingly, several members of the hnRNP (heterogenous nuclear ribonucleoprotein) family were identified (hnRNPA1, hnRNPA2B1, hnRNPK, hnRNPL and hnRNPU) as potential miR-128 targets and ORF1p interaction partners (Figure 1A). To validate the findings in the primary screen, we generated lentiviruses containing plasmids encoding miR-128, anti-miR-128 or scramble miR control. HeLa cells were transduced and selected with puromycin to generate stable miR-128, anti-miR-128 or miR control lines. We included analysis of TNPO1, which we have previously validated to be a miR-128 target (Idica et al, *in revisions*). The relative mRNA expression of hnRNPA1, hnRNPA2B1, hnRNPK, hnRNPL and hnRNPU was measured by qRT-PCR (Figure 1B, right panel). hnRNPA1 mRNA was significantly reduced in cells overexpressing miR-128 relative to miR controls (Figure 1B, top left panel). In cells where endogenous miR-128 was neutralized by anti-miR-128, hnRNPA1, hnRNPA2B1, hnRNPK and hnRNPU mRNA was significantly increased (Figure 1B). The finding that hnRNPA1 mRNA is significantly decreased or increased corresponding with overexpression or neutralization of miR-128 respectively, suggests that the effect of miR-128 on hnRNPA1 mRNA levels is specific. As hnRNPA1 plays an important role in nuclear transport by interacting with TNPO1 and is known to interact with ORF1p [42-50, 61], we decided to focus on the regulation of hnRNPA1 and examine if hnRNPA1 is required for miR-128-induced restriction of L1 mobilization.

**Figure 1:**
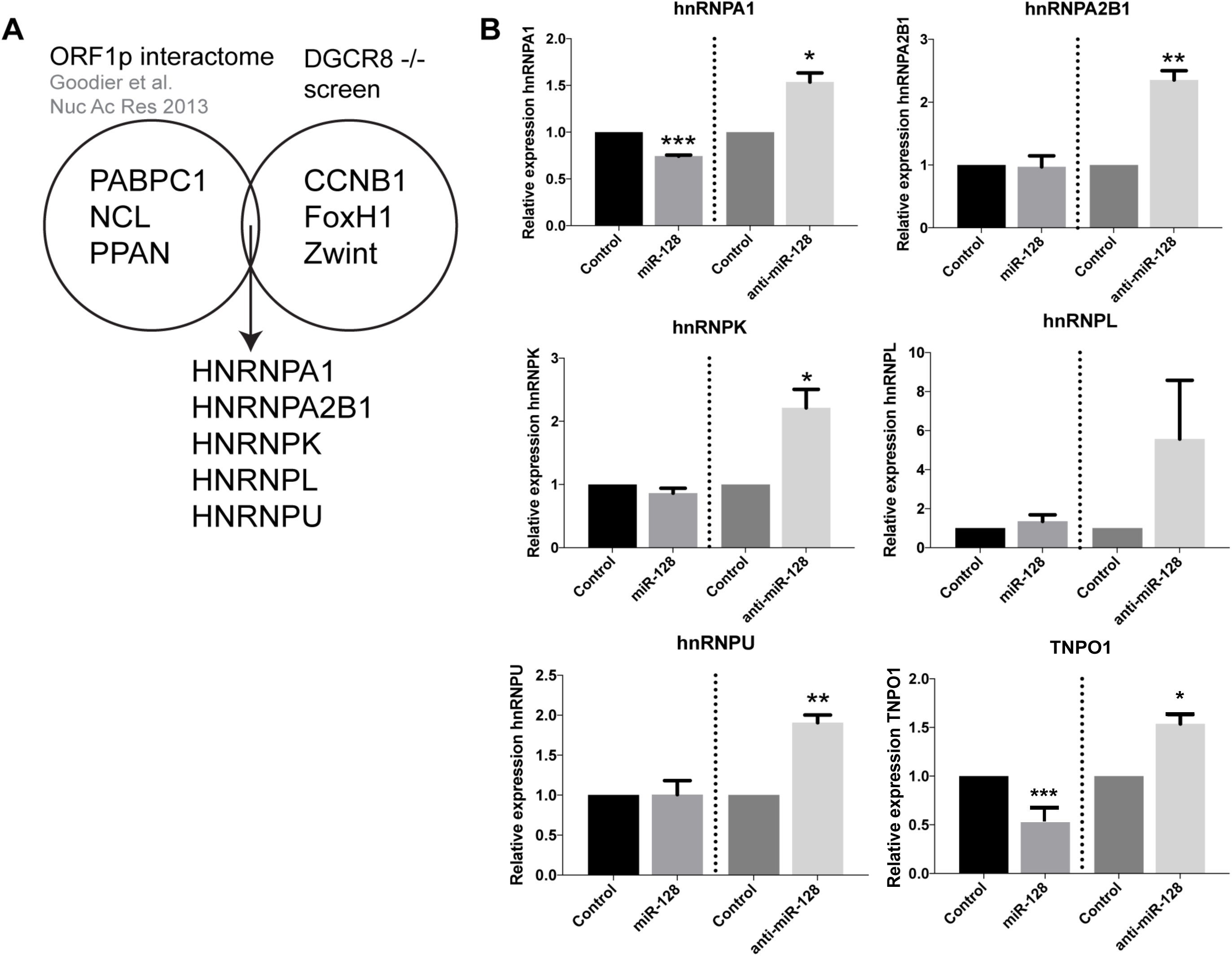
Identification of hnRNPA1 as a cellular target of miR-128. **(A)** Venn-diagram representing the L1 ORF1p interactome [50] and differentially expressed genes from the mESC DGCR8^-/-^ screen. Many members of the hnRNP family of proteins appeared in the intersection of these two datasets. **(B)** Relative amount of hnRNPA1, hnRNPA2B1, hnRNPK, hnRNPL and hnRNPU and TNPO1 (positive control) mRNA normalized to B2M determined in HeLa cells stably transduced with control miR, anti-miR-128 or miR-128 are shown as mean ± SD (n=3 technical replicates, *, p<0.05, **, p<0.01, ***, p<0.001).

### miR-128 reduces hnRNPA1 mRNA and protein levels

We next examined the effects of miR-128 on hnRNPA1 by performing and validating stable miR transductions with transient miR transduction of HeLa cells. We found that both transient and stable miR transduction of miR-128 resulted in significant reduced hnRNPA1 levels, and that miR-128 neutralization enhanced hnRNPA1 mRNA levels in both experimental conditions, relative to miR controls (Figure 2A). We further determined that miR-128 regulates hnRNPA1 in an induced pluripotent stem cell line, in a cancer initiating cell line and in three different cancer cell lines, and (iPSCs, colon cancer initiating cells (CCIC), breast cancer cell line (MDA-MB-231), non-small cell lung cancer line (NCI-A549) and a teratoma cell line (Tera-1). miR-128 significantly reduced hnRNPA1 in all but the lung cancer cell line and anti-miR-128 showed substantial enhanced hnRNPA1 levels in all cell lines except the Teratoma cell line (Figure 2B). Next we determined that miR-128 overexpressing HeLa cells showed significantly reduced hnRNPA1 protein levels and anti-miR-128 significantly enhanced hnRNPA1 protein amounts, relative to miR control HeLa cells, correlating with hnRNPA1 mRNA levels (Figure 2C). We finally validated that miR-128 regulates hnRNPA1 protein levels in additional 3 cancer cell lines (A549 (lung cancer), SW620 (colon cancer) and PANC1 (pancreatic cancer)). Together, these results show that miR-128 regulates hnRNPA1 mRNA and protein levels in multiple cell types.

**Figure 2:**
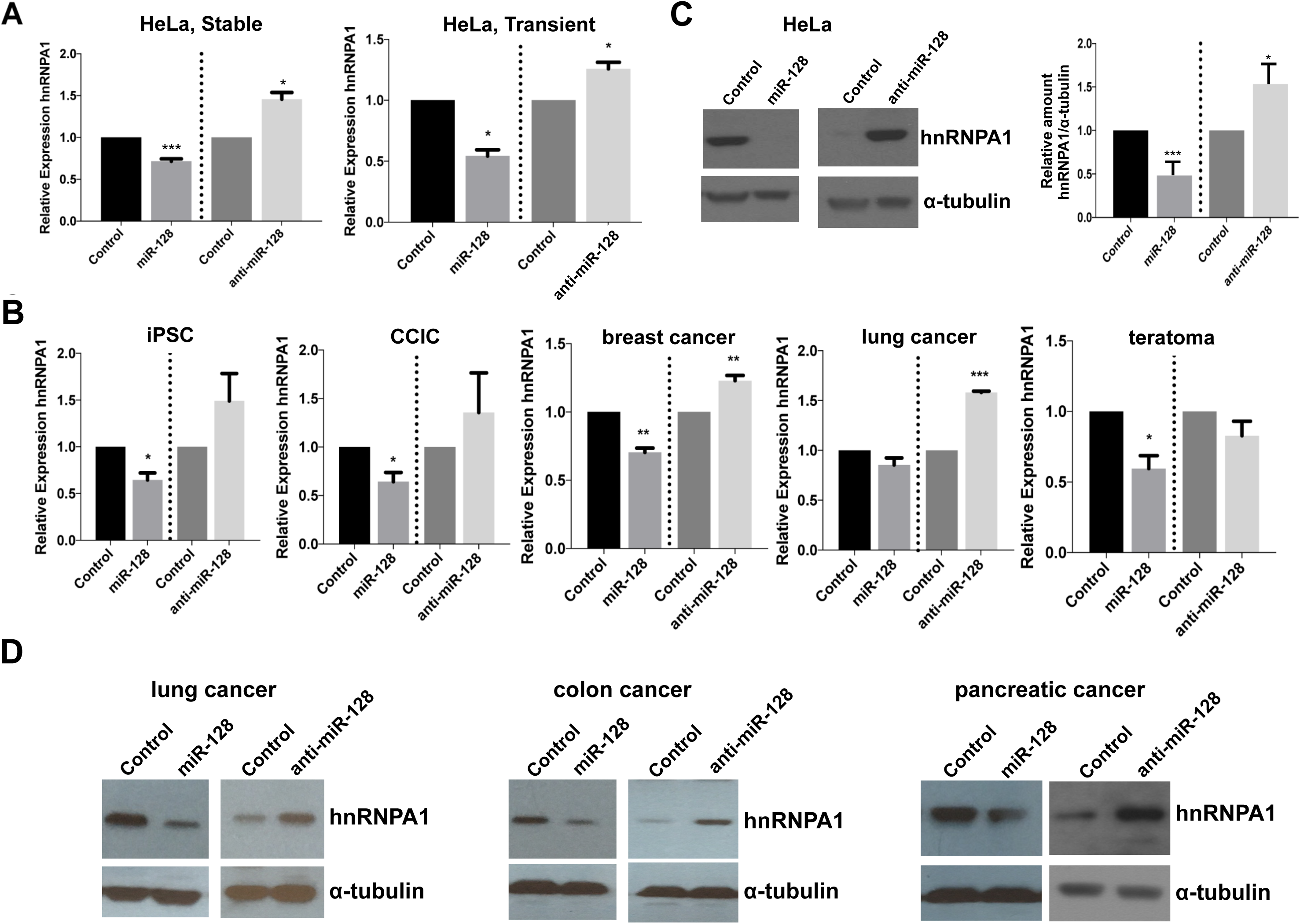
miR-128 reduces hnRNPA1 levels whereas miR-128 neutralization enhances hnRNPA1 levels in multiple cell types. **(A)** Relative amount of hnRNPA1 mRNA normalized to B2M in HeLa cells stably transduced with miR-128, anti-miR-128 or control constructs (left panel, n=2 independent biological replicates, p=ns); or transiently transfected with miR-128, anti-miR-128 or control mimics (right panel, mean ± SEM, n=3 independent biological replicates, *, p<0.05) **(B)** Relative amount of hnRNPA1 mRNA normalized to B2M in induced pluripotent stem cells, colorectal cancer initiating cells (CCIC), breast cancer cells (MDA-MB-231), non-small cell lung cancer (A549) cells and teratoma (Tera-1) cells **(C)** Immunoblot analysis of hnRNPA1 and α-tubulin protein levels in lysates from HeLa cells transduced with miR-128, anti-miR-128 or miR control constructs (left panel). Quantification of blots are shown (right panel) **(D)** Immunoblot analysis of hnRNPA1 and α-tubulin protein levels in protein-containing lysates isolated from non-small cell lung cancer (A549), colon cancer (SW620) (PANC1) cells transduced with miR-128, anti-miR-128 or miR control constructs. (n=3 independent biological replicates, mean ± SEM, *, p<0.05, **, p<0.01, ***, p<0.001).

### miR-128 binds directly to the CDS of hnRNPA1 mRNA

We next wished to determine if hnRNPA1 mRNA is a direct target of miR-128. When performing bioinformatics analyses of potential miR-128 binding sites in hnRNPA1 mRNA, we identified one potential 7-mer seed site in the coding DNA sequence (CDS) (Figure 3A). The CDS sequence of hnRNPA1 including the seed site was cloned into a miR-binding site luciferase reporter construct. As a positive control construct, a 23nt with perfect complementarity was generated as well. HeLa cells were co-transfected with the hnRNPA1 one of the binding site-encoding plasmid and either mature miR-128 or miR control mimics. Luciferase activity was significantly reduced in cells co-transfected with miR-128 and the plasmid encoding the hnRNPA1 binding site relative to those co-transfected with the miR control mimic and hnRNPA1 binding site plasmid (Figure 3B, left panel). These results indicate that miR-128 binds to the seed site located in the CDS of hnRNPA1. To further characterize the specificity of miR-128 binding to the seed site in hnRNPA1, a mutated seed site was generated and cloned into the miR-binding site luciferase reporter plasmid (Figure 3A). HeLa cells were co-transfected with one of the binding site-encoding plasmid (WT, seed mutant or positive control) and either mature miR-128 or miR control mimics. Luciferase activity was again significantly lower in HeLa cells transfected with miR-128 mimic and WT hnRNPA1, relative to miR control mimic suggesting that miR-151a can bind to the WT hnRNPA1 mRNA sequence and prevent the translation of luciferase (Figure 3B, left panel). In contrast, HeLa cells transfected with miR-128 mimic and the mutant hnRNPA1 CDS site, exhibited de-repressed luciferase activity to the same levels as the WT hnRNPA1 and miR-control cells; consistent with the conclusion that miR-128 no longer binds and represses reporter gene expression (Figure 3B, right panel).

**Figure 3:**
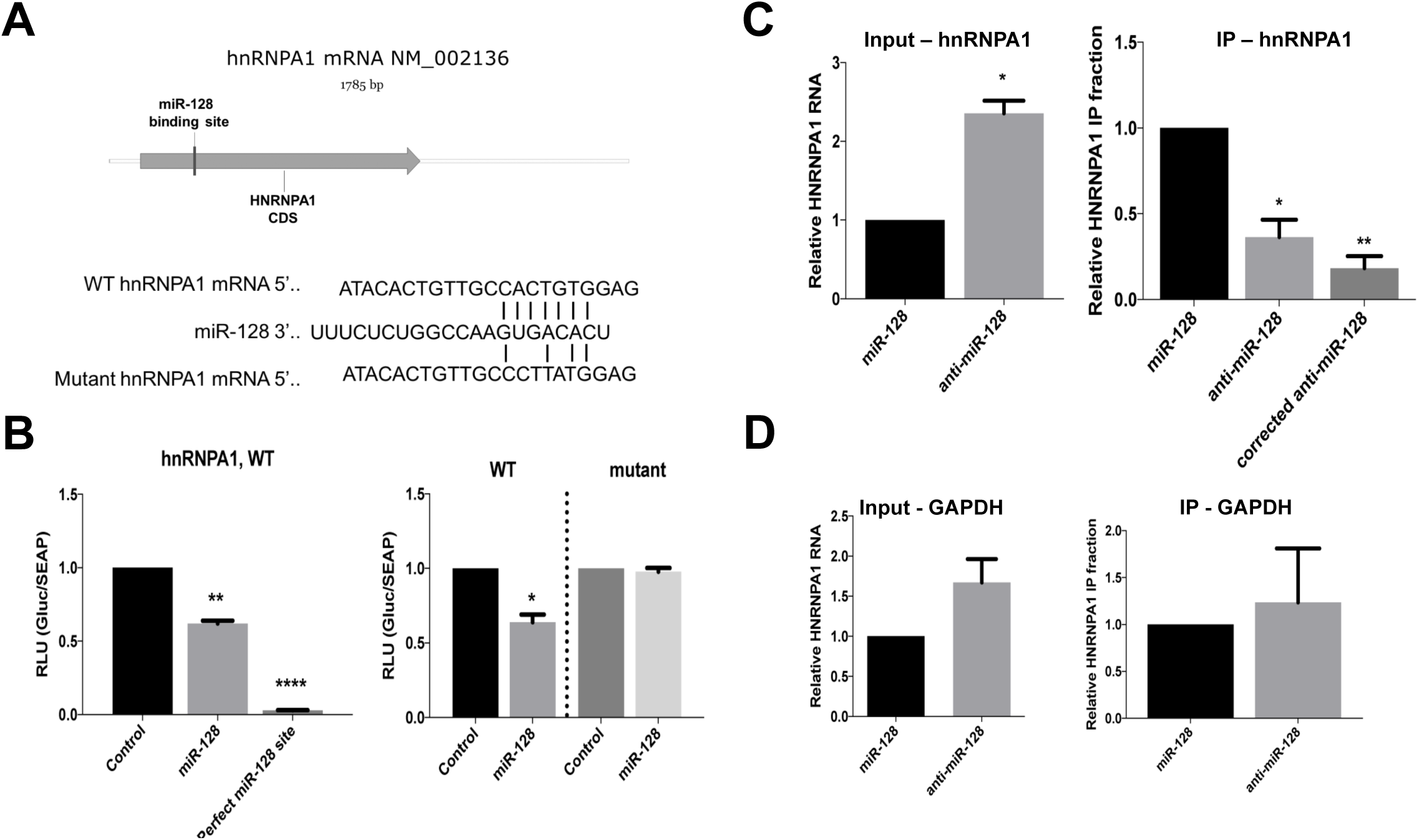
miR-128 represses hnRNPA1 expression by binding directly to CDS RNA. **(A)** Schematic of the predicted miR-128 7-mer binding site in the coding region (CDS) of hnRNPA1 mRNA (top panel). Predicted base pairing of miR-128 to the seed sequence of wild-type (WT) hnRNPA1 as well as a representation of mutations in the seed sequence (mutant) used for luciferase binding assays. **(B)** Relative luciferase activity in HeLa cells transfected with plasmids expressing a Gaussia luciferase gene fused to the wild-type (WT) binding site or positive control sequence corresponding to the 22 nucleotide perfect match of miR-128 and co-transfected with control or mature miR-128 mimics were determined 48 hours post-transfection (left panel, n=3 independent biological replicates, mean ± SEM, **, p<0.01, ****, p<0.0001). Relative luciferase activity in HeLa cells transfected with plasmids expressing the luciferase gene fused to the WT or mutated binding site (mutant) and co-transfected with control or mature miR-128 mimics were determined 48 hours post-transfection (right panel, n=3 independent biological replicates, mean ± SEM, *, p<0.05). **(C)** Argonaute-RNA immuno-purification in HeLa cell lines stably transduced with miR-128 overexpression or miR-128 neutralization (anti-miR-128) was performed. Relative amounts of hnRNPA1 RNA normalized to B2M was determined for input samples (top left panel “input – hnRNPA1”, n=3 independent biological replicates, mean ± SEM, *, p<0.05). Relative fraction of hnRNPA1 transcript amounts associated with immune-purified Ago complexes is shown for IP samples, hnRNPA1 fractions normalized to the amount of TNPO1 in input are shown as “corrected” (top right panel “IP – hnRNPA1”, n=3 independent biological replicates, mean ± SEM, *, p<0.05, **, p<0.01). **(D)** Relative amount of GAPDH in the same input and IP samples were determined as a negative control. (top right panel “IP – hnRNPA1”, n=3 independent biological replicates, mean ± SEM).

Finally, in order to evaluate miR-128 directly interacts with hnRNPA1 mRNA in cells, Argonaute-RNA immuno-purification (RIP) was performed. Briefly, transduced HeLa cell lines stably expressing miR-128 or anti-miR-128 were lysed and Argonaute complexes relatively enriched with miR-128 (miR-128) or depleted of miR-128 (anti-miR-128) were isolated. The argonaute complexes were then analyzed for occupancy by hnRNPA1 mRNA by qPCR; if hnRNPA1 mRNA is a direct target of miR-128, occupancy should be higher in complexes enriched with miR-128 relative to those depleted of miR-128 (anti-miR-128). As expected, the relative level of hnRNPA1 mRNA was significantly lower in cells stably overexpressing miR-128 compared to cells expressing anti-miR-128 (Figure 3C, top left panel “Input”). The relative fraction of Argonaute-bound hnRNPA1 mRNA was significantly increased in cells overexpressing miR-128 relative to cells expressing anti-miR-128 (Figure 3C, top right panel “IP”). When correcting for the higher amount of hnRNPA1 mRNA in anti-miR-128 input samples, the relative fraction of Argonaute-bound hnRNPA1 mRNA in IP samples were found to be more significantly reduced, relative to miR-128 samples (Figure 3C, top right panel “corrected”). miR-128 did not reduce GAPDH mRNA amounts or immuno-purified GAPDH mRNA (Figure 3D). These data combined, suggests that induced amounts of miR-128 result in enrichment of hnRNPA1 mRNA bound to Argonaute complexes by a direct interaction with the seed sequence in the CDS of hnRNPA1 mRNA.

### L1 retrotransposition is dependent on hnRNPA1

hnRNPA1 is a known binding partner of L1 ORF1p [50] and TNPO1 (Idica et al. *in revisions*) suggesting a possible role for hnRNPA1 in the L1 retrotransposition life cycle. We first wished to determine whether hnRNPA1 is required for successful *de novo* retrotransposition of L1 and secondly whether miR-128-induced L1 restriction is dependent on reduced hnRNPA1 levels.

We performed colony formation assays using different variants of a neomycin reporter constructs encoding the full length L1 mRNA and a retrotransposition indicator cassette. The WT L1 construct consists of a neomycin gene in the antisense orientation relative to a full-length L1 element, which is disrupted by an intron in the sense orientation. In order for the neomycin (neo) protein to be translated into a functional enzyme, L1 transcription and splicing of the mRNA, reverse transcription followed by integration of the spliced variant into the genome, is required. This assay therefore allows for the quantification of cells with new (*de novo*) retrotransposition and genomic integrations events in culture. In addition, we have generated a miR-128 resistant variant of the L1 plasmid, by introducing a silent mutation in the miR-128 binding site (in the ORF2 sequence) attenuating miR-128 binding, but allowing L1 to retrotranspose (as described in [32] and Idica et al. *in revisions*). Finally a third variant of the L1 plasmid described by [62] encodes a L1 RNA containing a D702A mutation in the reverse transcriptase (RT) domain of the ORF2 protein, rendering the encoded L1 RT deficient (RT dead). This plasmid variant was used as a negative control as previously described (Idica et al., *in revisions*). We generated stable HeLa cell lines overexpressing full length hnRNPA1 or plasmid control, shRNA against hnRNPA1 or a control sequence (GFP, Green Fluorescent Protein). Based on our recent finding that TNPO1 is required for L1 to successfully transpose and that miR-128 targets TNPO1 mRNA for degradation, we also wished to examine the potential additive effect of TNPO1 and hnRNPA1 on L1 retrotransposition, both being miR-128 targets. For this reason we also generated HeLa cell lines stably overexpressing both hnRNPA1 and TNPO1 (hnRNPA1 + TNPO1) and cell lines overexpressing both sh-hnRNPA1 and sh-TNPO1 (sh-hnRNPA1 + and sh-TNPO1). We validated that hnRNPA1 and TNPO1 was regulated as expected in the cell lines (Figure 4 and data now shown) and then initiated colony formation assay analysis. All hnRNPA1/TNPO1 modulated HeLa cell lines were transfected with the WT L1 or the RT deficient L1 (RT-dead) neomycin reporter and selected for 14 days with neomycin replenished daily. We observed a significant decrease in neomycin resistant colonies in cells overexpressing sh-hnRNPA1, relative to control HeLa cells (Figure 4A). Double knock-down (sh-hnRNPA1 and sh-TNPO1) HeLa cells showed further reduced levels of L1 retrotransposition (Figure 4A). In contrast, cells overexpressing hnRNPA1 were characterized by a significantly increased number of neo resistant colonies, compared to control HeLa cells (Figure 4B). The increase in neo resistant colonies was further increased in hnRNPA1 + TNPO1 double over-expressing cells (Figure 4B). Importantly, hnRNPA1 and TNPO1 modulated HeLa cells encoding RT-deficient L1 (RT-dead) resulted in no neomycin-resistant colonies, demonstrating that colonies obtained upon wild-type L1 plasmid transfections and neo selection are the consequence of a round of *de novo* L1. These experiments demonstrate that hnRNPA1 is required for optimal L1 activity.

**Figure 4:**
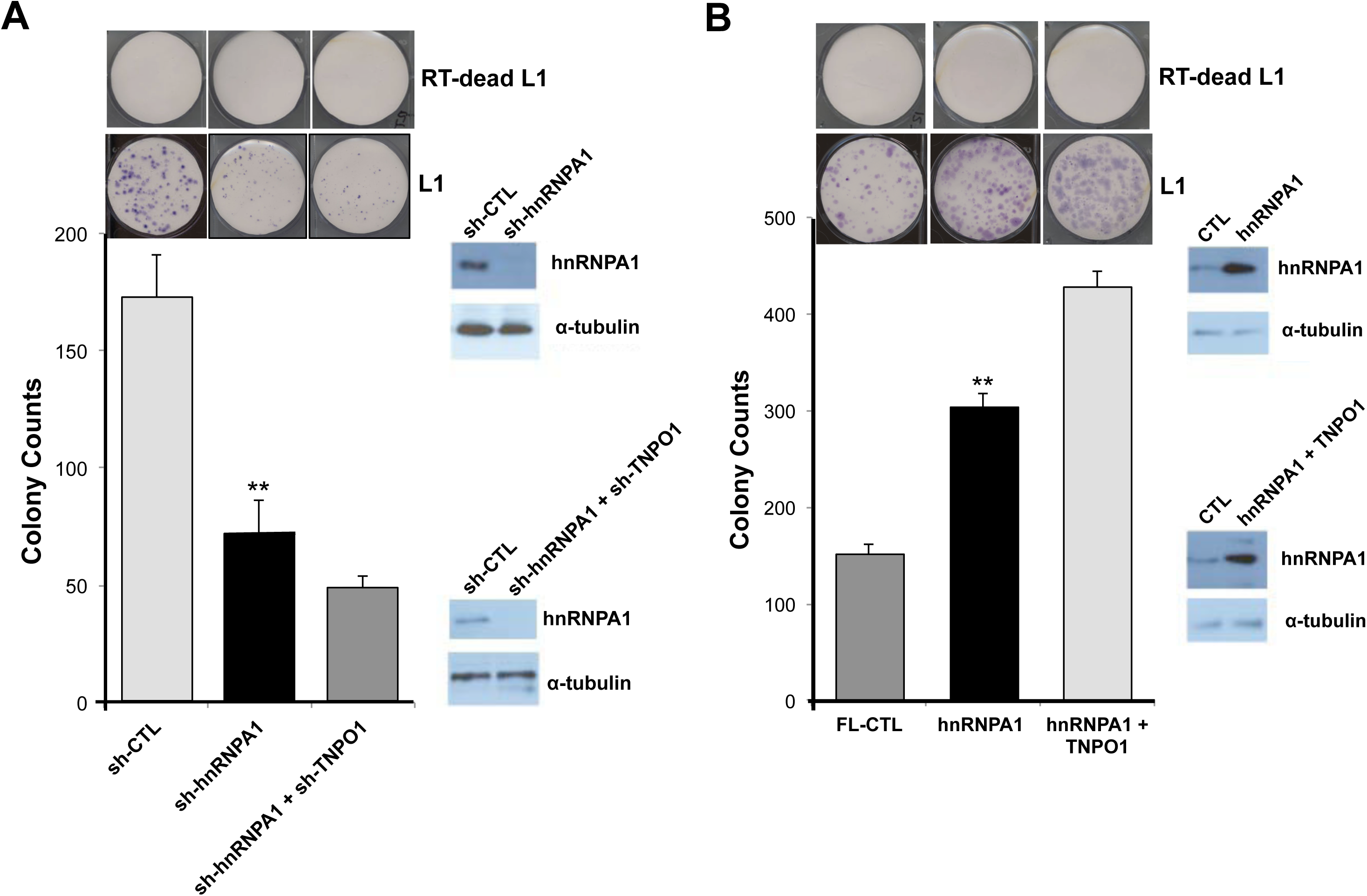
hnRNPA1 knockdown reduces *de novo* L1 retrotransposition and hnRNPA1 induction enhances L1 mobilization. **(A)** Representative example of neomycin-resistant colony counts from colony formation assay in HeLa cells stably transduced with plasmids encoding shRNA against GFP (control), hnRNPA1 or shRNPA1 and TNPO1 and transfected with WT L1 (L1) or RT deficient L1 (RT-dead L1) expression construct. (n=3 technical replicates, mean ± SD, ***, p<0.001). WB analysis validates reduced levels of hnRNPA1 in knock-down cell lines **(B)** *De novo* retrotransposition was determined by colony formation assay in HeLa cells stably transduced with plasmids encoding control plasmid (control), hnRNPA1 or hnRNPA1 and TNPO1 and transfected with L1 expression construct. WB analysis validates modulated levels of hnRNPA1 in knock-down and over-expressing cell lines. (n=3 technical replicates, mean ± SD, *, p<0.05, **, p<0.01),

### miR-128-induced L1 restriction is partly dependent on hnRNPA1

At this point our findings show that L1-mobolization is dependent on repressing the cellular co-factor hnRNPA1, TNPO1, and direct binding to L1 RNA. In order to evaluate the relative importance of miR-128-induced hnRNPA1 repression we next performed rescue experiments. In brief, we generated miR-128 or miR control HeLa cell lines in which we co-expressed either full lenght hnRNPA1 or plasmid controls. All HeLa cell lines were then transfected with the mutant L1 (miR-128 resistant) plasmid or the RT-dead L1 plasmid as previously described (Idica et al. *in revisions*). As expected miR-128 significantly reduced L1 retrotransposition as determined by reduced neo resistant colonies (Figure 5A). When reintroducing hnRNPA1 into miR-128 over-expressing HeLa cells, we observed a partial, but significant rescue of miR-128 induced L1 restriction (Figure 5A), whereas all HeLa cell lines transfected with the RT dead L1 plasmid, resulted in no colonies following neo selection.

**Figure 5:**
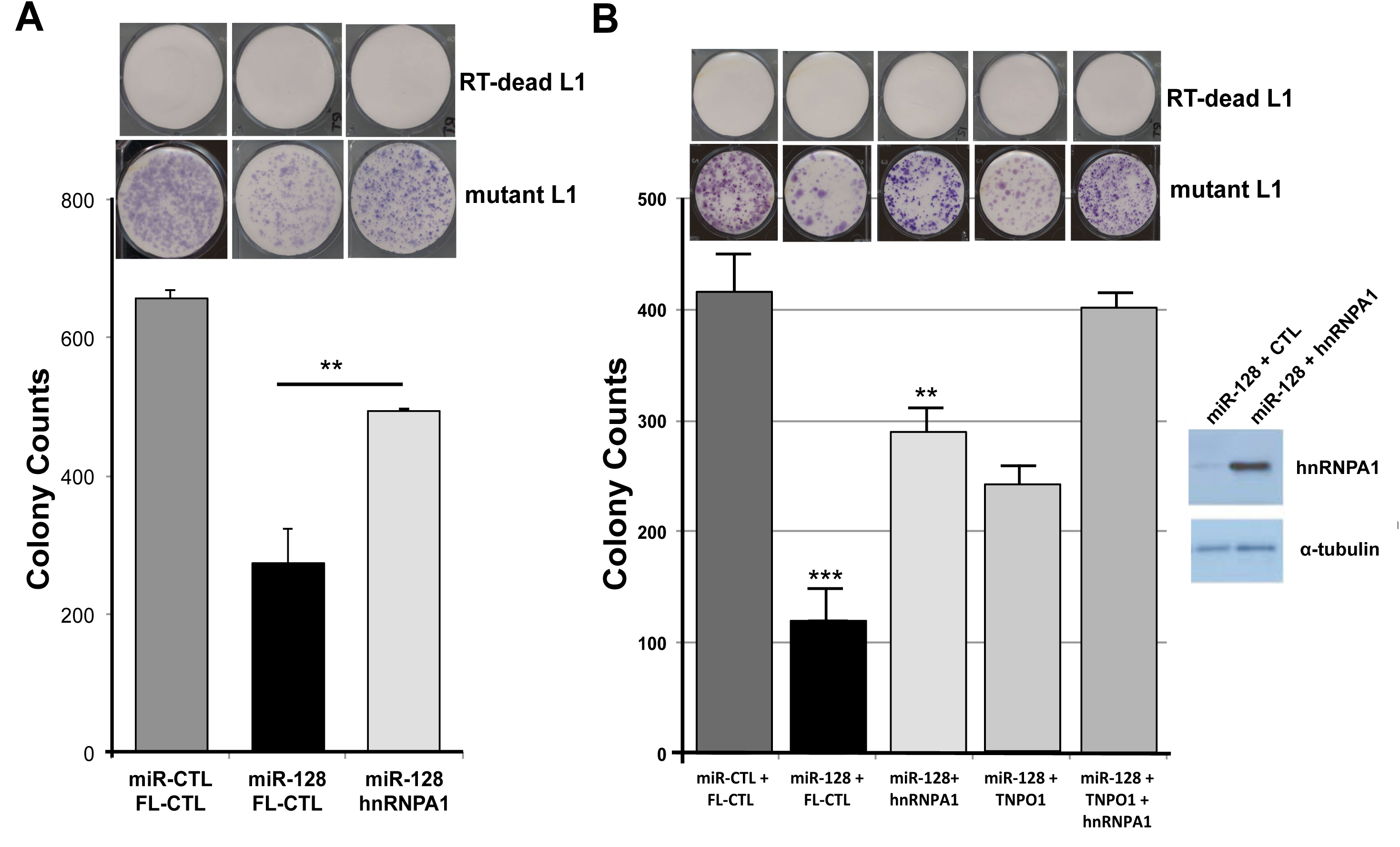
hnRNPA1 partly rescues miR-128-induced inhibition of *de novo* L1 retrotransposition. **(A)** The functional importance of hnRNPA1 in miR-128-induced L1 repression was evaluated by colony formation assays using mutant L1 (miR-128 resistant) or RT deficient L1 (RT-dead L1) expression constructs in stable HeLa cell lines expressing either miR-control (miR-CTL) or miR-128 along with either a plasmid control (FL-CTL) or induced hnRNPA1 (hnRNPA1) **(B)** Colony formation assays was performed mutant L1 (miR-128 resistant) or RT deficient L1 (RT-dead L1) expression constructs in stable HeLa cell lines expressing either miR-control (miR-CTL) or miR-128 along with either a plasmid control (FL-CTL) or induced hnRNPA1 (hnRNPA1) or TNPO1 (TNPO1) or both hnRNPA1 and TNPO1 (hnRNPA1 + TNPO1) (double rescue). (n=3 technical replicates, mean ± SD, **, p<0.01). WB analysis validates increased levels of hnRNPA1 in hnRNPA1 rescue cell line.

Finally, we performed a double rescue experiment using miR control and miR-128 HeLa cells, which overexpressed either plasmid control, full length hnRNPA1, full length TNPO1 or both hnRNPA1 + TNPO1. As observed previously, miR-128 significantly reduced L1 activity, and hnRNPA1 partly rescue the inhibitory effect of miR-128, as compared to control cells (Figure 5B). In addition, as previously shown TNPO1 can also partly rescue *de novo* retrotransposition, and the potency of rescue was further enhanced in cell overexpressing both hnRNPA1 and TNPO1, relative to plasmid control (Figure 5B and Idica et al. *in revisions*). These results support the idea that miR-128 functions through direct binding of L1 RNA and by regulating at least two cellular co-factors (hnRNPA1 and TNPO1), which L1 is dependent on for successful mobilization.

## DISCUSSION

Interactions between hnRNPA1 and TNPO1, as well as hnRNPA1 and L1 ORF1p have been reported [42-48, 50, 61], however no direct role for hnRNPA1 on L1 retrotransposition has been identified. Our results demonstrate that miR-128 functions to repress hnRNPA1 as one of the cellular co-factors in the L1 retrotransposition pathway and complement our earlier work describing miR-128-induced direct repression of L1 retrotransposons [32] and miR-128-induced repression of a nuclear import factor of L1 (TNPO1) (Idica et al. *in revisions*). This body of work agrees with the concept that miRs repress multiple cellular targets in unison to regulate important cellular pathways [33-36].

Our findings also support the conclusion that miR-128 significantly reduces hnRNPA1 levels by directly interacting with a miR-128 seed site in the coding region sequence (CDS) of hnRNPA1 mRNA. The finding that miR-128 targets the CDS of hnRNPA1 resulting in significantly reduced hnRNPA1 mRNA and in particular hnRNPA1 protein levels in a small panel of different cell types, was surprising. However, it is well established that miRs regulate gene products by preferentially interacting with the 3’UTR and/or the CDS of target mRNA and that such interactions are of functional importance [32, 63, 64]. In addition, we have demonstrated for the first time that *do novo* retrotransposition of L1 elements are dependent on hnRNPA1 (by overexpression and shRNA experiments), and that hnRNPA1 is important for miR-128-induced inhibition of L1 activity, as overexpression of hnRNPA1 in miR-128 HeLa cells partially can rescue the effect of miR-128 on L1 replication. Finally, double knock-down, double over-expression and double rescue experiments of hnRNPA1 and TNPO1 indicate that both proteins are involved in the L1 life cycle. Additional studies are needed to determine whether additional co-factors required by L1, are regulated by miR-128. Mechanistic studies are also needed to dissect what exact role hnRNPA1 plays in L1 retrotransposition. It is likely that hnRNPA1 and TNPO1 corporate at facilitating active nuclear import of some L1-RNP complexes, and thus our finding that miR-128 target both mRNAs support a possible role for miR-128 in determining whether L1-RNP complexes gain access to host DNA, independently of cell division. hnRNPA1 is also involved in many other processes necessary for cellular function including proliferation [65], mRNA splicing [66], and telomerase activity [67], and mutations in hnRNPA1 have been associated with human diseases including amyotrophic lateral sclerosis 20 [68], lung adenocarcinoma [69] and HIV-1 [70, 71].

In summary, we have identified hnRNPA1 as a novel miR-128 target. We have determined that miR-128 significantly reduces hnRNPA1 protein and mRNA amounts by directly interacting with the coding sequence of hnRNPA1 mRNA. In addition we have demonstrated that hnRNPA1 is required for L1 retrotransposition and for miR-128-induced L1 repression. This body of work suggests that miR-128 represses L1-induced mutagenesis through a multi-facetted mechanism by both directly targeting of L1 RNA and indirectly through the repression of cellular co-factors, which L1 is dependent on (TNPO1 and hnRNPA1) ([32] and Idica et al. *in revisions*).

## ACKNOWLEDGEMENTS

We thank S. Sardari for lab assistance in quantifying results. We also thank J. Moran (University of Michigan Medical School, Ann Arbor, MI) and M. An (Washington State University, Pullman, WA) for generously sharing pJM101/L1 and pWA355 plasmids. This work was supported by University of California Cancer Research Coordinating Committee 55205 (I.M.P.), American Cancer Society – Institutional Research Grant 98-279-08 (I.M.P.), University of California Irvine Institute for Memory Impairments and Neurological Disorders grant (I.M.P.).

## AUTHOR CONTRIBUTION

L. Fung made the initial identification of hnRNPA1 as a potential miR-128 target and an ORF1 interaction partner. E. Sevrioukov, H. Guzman and A. Idica performed the majority of experiments, with the help of D. Jury and I. Daugaard, demonstrating that miR-128 targets hnRNPA1 and that miR-128-induced L1 restriction is partly dependent on hnRNPA1. A. Mortazavi and E. Park was responsible for RNA sequencing and data analysis. D. Zisoulis performed the Ago RNA IPs, I. Daugaard also helped generate the final figures and edited the manuscript and IM. Pedersen performed some experiments, directed all experiments, figure design and wrote the manuscript.

## MATERIALS AND METHODS

### Cell culture

All cells were cultured at 37°C and 5% CO2 and routinely checked for mycoplasma (LT07-218, Lonza). HeLa cells (CCL-2, ATCC) were cultured in EMEM (SH3024401, Hyclone) supplemented with 10% HI-FBS (FB-02, Omega Scientific), 5% Glutamax (35050-061, ThermoFisher), 3% HEPES (15630-080, ThermoFisher), and 1% Normocin (ant-nr-1, Invivogen). 293T cells (CRL-3216, ATCC) used to generate lentiviruses, H23 cells (CRL-5800, ATCC), and MDA-MB-231 (ATCC HTB-26) were cultured in DMEM supplemented with 10% HI-FBS (FB-02, Omega Scientific), 5% Glutamax (35050-061, Lifetech) and 1% Normocin (ant-nr-1, Invivogen). Tera-1 cells (HTB-105, ATCC) were cultured in McCoy’s 5A (16600-082, Lifetech) supplemented with 20% Cosmic Serum (SH3008702, Fisher Sci), and 1% Normocin (ant-nr-1, Invivogen). Passaging was performed at 80% confluence with 0.25% trypsin (SH30042.01, Hyclone). Colon cancer initiating cells (CCICs, gifted from Professor Marian Waterman, UC Irvine) were verified and cultured as described in Sikandar et al (Sikandar et al., n.d.). Briefly, CCICs were cultured as spheres in ultra-low attachment flasks in DMEM/F12, N2 supplement (17502-048, Lifetech), B27 supplement (17504-044, Lifetech), heparin (4 μg/mL, Sigma), epidermal growth factor (20 ng/mL), and basic fibroblast growth factor (20ng/mL). H23 (CRL-5800, ATCC) were cultured in RPMI-1640 (11875, Lifetech), 10% HI-FBS, 5% Glutamax, and 1% Normocin. mESCs were maintained in knockout DMEM medium (Invitrogen) supplemented with 15% fetal bovine serum, LIF and 2i (PD0325901 and CHIR99021) as per standard techniques.

### Transfection and transduction

OptiMem (31985070, Lifetech) and Lipofectamine RNAiMAX (13778075, Lifetech) were used to complex and transfect 20μM miR-128 mimic, anti-miR-128 or control mimics (C-301072-01 and IH-301072-02, Dharmacon) into cells. OptiMem and Xtreme Gene HP (06366236001, Roche Lifescience) was used to transfect pJM101 neomycin L1 reporter plasmid into HeLa cells. Cells were transduced with high titer virus using polybrene (sc-134220, Santa Cruz Biotech) and spinoculation (800xg at 32°C for 30 minutes). Transduced cells were then selected and maintained using 3μg/mL puromycin.

### RNAi using shRNA against hnRNPA1

shRNA for hnRNPA1 was designed using the RNAi Consortium (https://www.broadinstitute.org/rnai/public/) using clone TRCN0000235098 and cloned into pLKO.1 puro backbone (Addgene, #8453). pLKO shGFP control plasmid was pre-assembled (Addgene, #30323).

### Lentiviral packaging

VSVG-pseudotyped lentiviral vectors were made by transfecting 0.67μg of pMD2-G (12259, Addgene), 1.297μg of pCMV-DR8.74 (8455, Addgene), and 2μg of mZIP-miR-128, mZIP-anti-miR-128, pLKO-shControl or pLKO-shHNRNPA1 (transfer plasmid)) into 293T cells using Lipofectamine LTX with plus reagent (15338030, ThermoFisher). Virus-containing supernatant was collected 48hr and 96hr post-transfection. Viral SUPs were concentrated using PEG-it virus precipitation solution (LV810A-1) according to manufacturer’s instructions.

### RNA extraction and quantification

RNA was extracted using Trizol (15596-018, ThermoFisher) and Direct-zol RNA isolation kit (R2070, Zymo Research). cDNA was made with High-Capacity cDNA Reverse Transcription Kit (4368813, ThermoFisher). Amount of hnRNPA1 mRNA was analyzed by qRT-PCR (Sense primer 5’-aagcaattttggaggtggtg-3’; Antisense primer 5’-atagccaccttggtttcgtg-3’) using Forget-me-not qPCR mastermix (Biotium) relative to beta-2-microglobulin (B2m, Sense primer 5’-ATGTCTCGCTCCGTGGCCTTAGCT-3’; Antisense primer 5’-TGGTTCACACGGCAGGCATACTCAT-3’) housekeeping gene and processed using the δδC_t_ method.

### Western blotting

Rabbit anti-human hnRNPA1 antibody (K350, Cell Signaling Technology) was used at 1:2000. Anti-alpha Tubulin antibody (ab4074, Abcam) was diluted 1:5000 and used as a loading control, validation of antibodies can be found on the manufacturer websites. Secondary HRP-conjugated anti-rat (ab102172, Abcam) or HRP-conjugated anti-rabbit (#NA934, GE lifesciences) were used at 1:5000. ECL substrate (32106, ThermoFisher) was added and visualized on a BioRad ChemiDoc imager.

### Argonaute-RNA immuno-purification

Immunopurification of Argonaute from HeLa cell extracts was performed using the 4F9 antibody (#sc-53521, Santa Cruz Biotechnology) as described previously [72, 73]. Briefly, 10mm plates of 80% confluent cultured cells were washed with buffer A [20 mM Tris-HCl pH 8.0, 140 mM KCl and 5 mM EDTA] and lysed in 200ul of buffer 2XB [40 mM Tris-HCl pH 8.0, 280 mM KCl, 10 mM EDTA, 1% NP-40, 0.2% Deoxycholate, 2X Halt protease inhibitor cocktail (Pierce), 200 U/ml RNaseout (Life Technologies) and 1 mM DTT. Protein concentration was adjusted across samples with buffer B [20 mM Tris-HCl pH 8.0, 140 mM KCl, 5 mM EDTA pH 8.0, 0.5% NP-40, 0.1% deoxycholate, 100 U/ml Rnaseout (Life Technologies), 1 mM DTT and 1X Halt protease inhibitor cocktail (Pierce)]. Lysates were centrifuged at 16,000g for 15 mins at 4°C and supernatants were incubated with 10-20 ug of 4F9 antibody conjugated to epoxy magnetic beads (M-270 Dynalbeads, Life Technologies) for 2 hours at 4°C with gentle rotation (Nutator). The beads, following magnetic separation, were washed three times five mins with 2 ml of buffer C [20 mM Tris-HCl pH 8.0, 140 mM KCl, 5 mM EDTA pH 8.0, 40 U/ml Rnaseout (Life Technologies), 1 mM DTT and 1X Halt protease inhibitor cocktail (Pierce). Following immunopurification, RNA was extracted using miRNeasy kits (QIAGEN), following the manufacturer’s recommendations and qPCR was performed using hn RNPA1 primers designed around the binding site of miR-128 (Sense primer 5’-TCTCCTAAAGAGCCCGAACA-3’; Antisense primer 5’-TTGCATTCATAGCTGCATCC-3’) or GAPDH (Sense primer 5’-GGTGGTCTCCTCTGACTTCAA-3’; Antisense primer 5’-GTTGCTGTAGCCAAATTCGTT-3’) normalized to B2m (Sense primer 5’-ATGTCTCGCTCCGTGGCCTTAGCT-3’; Antisense primer 5’-TGGTTCACACGGCAGGCATACTCAT-3’). Results were normalized to their inputs.

### Cloning

To generate the hnRNPA1 and TNPO1 full-length clones we modified the plasmid pFC-PGK-MCS-pA-EF1-GFP-T2A-Puro (SBI, backbone) by replacing the PGK with a CMV promoter. The CMV promoter provides a strong and robust expression on most cell types. The CMV promoter was amplified by PCR from the phiC31 integrase expression plasmid (SBI). To generate the CMV promoter insert, we used the sense CMV primer (5’-CTAGAACTAG TTATTAATAG TAATCAATTA CGGGGTC-3’) and antisense CMV primer (5’-GATATCGGAT CCACCGGTAC CAAGCTTAAG TTTAAAC-3’). The insert and the backbone of the plasmid were cut by *Xba*I and *BamH*I and purified by an agarose gel. Insert and backbone were ligated together using the quick ligation kit (NEB) and transformed. The resulting plasmid pFC-CMV-MCS-pA-EF-1-GFP-T2A-Puro-MH1 was verified by sequencing.

For the cloning of the full-length hnRNPA1 and TNPO1 mRNA expression clone, we isolated total RNA from HeLa cells. 20 ng of the total RNA was reverse transcribed using a poly dT primer. All amplicons were generated using the Phusion High-Fidelity PCR Kit (NEB) according to the manufacturer’s protocol. The fragments were stepwise assembled by using the cold fusion kit (SBI) and cloned into the pFC-CMV-MCS-pA-EF-1-GFP-T2A-Puro-MH1 *Bam*HI/*Cla*I linearized backbone by cold fusion. The resulting plasmid pFC-CMV-TNPO1-pA-EF-1-GFP-T2A-Puro-MH1 was verified by sequencing. FL-Control is an empty vector.

### Luciferase binding assay

Wild-type hnRNPA1 sense primer (5’-AATTCTTGGGTTTGTCACATATGCCACTGTGGAGGAGGTGGATGCAGCTA-3’) and antisense primer (5’-CTAGTAGCTGCATCCACCTCCTCCACAGTGGCATATGTGACAAACCCAA-3’), mutated hnRNPA1 sense primer (5’-AATTCTTGGGTTTGTCACATATGCCCTTATGGAGGAGGTGGATGCAGCTA-3’) and antisense primer (5’-CTAGTAGCTGCATCCACCTCCTCCATAAGGGCATATGTGACAAACCCAA-3’), or positive control sense primer (5’-AATTCAAAGAGACCGGTTCACTGTGAA-3’) and antisense primer (5’-CTAGTTCACAGTGAACCGGTCTCTTTG-3’) sequences were cloned into dual-luciferase reporter plasmid (pEZX-MT05, Genecopoeia). 3x10^5^ HeLa cells were forward transfected with 0.8μg of reporter plasmid (WT, mutated, Pos) and 20nM miR-128 mimic (Dharmacon) or Control mimic (Dharmacon) using Attractene transfection reagent (301005, Qiagen) according to manufacturer instructions. Relative Gaussia Luciferase and SEAP was determined using Secrete-Pair Dual Luminescence Assay Kit (SPDA-D010, Genecopoeia). Luminescence was detected by Tecan Infinite F200 Pro microplate reader.

### Site directed mutagenesis

Reverse transcriptase incompetent PJM101/L1 plasmid was made using Q5 Site-directed mutagenesis Kit (E0554S, New England Biolabs) and mutation strategy described in Morrish et al. where D702A mutation in L1 ORF2 resulted in an incompetent reverse transcriptase, called RT-dead L1.

### Colony formation assay

Stable HeLa lines expressing miR-128, anti-miR-128, scramble control, shControl, shhnRNPA1, shTNPO1, Full length hnRNPA1 or TNPO1, or plasmid control were transfected with pJM101/L1RP or RT-dead L1 plasmid (containing neomycin resistance retrotransposition indicator cassette) per well using X-treme gene HP DNA transfection reagent (06366236001, Roche) according to manufacturer instructions. Cells were selected using 500μg/mL G418 (ant-gn-1, Invivogen). Neomycin-resistant colonies were fixed with cold 1:1 methanol:acetone then visualized using May-Grunwald (ES-3410, Fisher Sci) and Jenner-Giemsa staining kits (ES-8150, Fisher Sci) according to manufacturer protocol. Stable iPSC lines expressing miR-128, anti-miR-128 or scramble control were transfected with pJM101/L1RP using Xtreme gene HP DNA transfection reagent according to manufacturer instructions. Selection began with 25 μg/mL G418 72hr post-transfection and selection was maintained with daily media changes until negative control (non-transfected) cells died. Neomycin-resistant colonies were fixed as described above.

### RNA sequencing and data analysis

DGCR8^-/-^ mESCs were transfected with miR controls or miR-128 in triplicate cultures and harvested cells after 12hours in order to enrich for primary target mRNAs, as opposed to studying secondary effects of miR-128. Two replicates of each triplicate were selected and cDNA libraries were generated using the Smart-seq2 protocol [57]. The libraries were sequenced as 43 bp paired-end reads. STAR [58] was used to align the reads on to the mm9 genome. RSEM [59] was used to quantitate the gene expression and EBSeq [60] was used to identify differentially expressed genes.

## Notes

**Conflict of interest:** The authors declare no conflict of interest.

